# Murine model of antibiotic-associated *S. aureus* gastrointestinal infections (SAGII) and colonization

**DOI:** 10.1101/2025.04.30.651547

**Authors:** Maria Niamba, Stephanie Yang, Liahm Blank, Liliko Watanabe, Efren Heredia, Ernesto Abel-Santos

## Abstract

**Background:** *Staphylococcus aureus* is an opportunistic pathogen that can both colonize the gastrointestinal tract and cause antibiotic associated diarrhea.

**Methods:** To develop a robust murine model for *S. aureus* gastrointestinal infection (SAGII) and colonization, mice were (a) treated with varying antibiotic regimes prior to infection, (b) infected with either a methicillin-sensitive *S. aureus* (MSSA) or a methicillin-resistant *S. aureus* (MRSA) strain, (c) challenged with different bacterial inocula, (d) tested for sexual dimorphism of SAGII virulence, and (e) tested for macronutrient effects on SAGII onset and virulence.

**Results:** We found that antibiotic-treated male mice (but not female mice) were highly susceptible to even low inoculums of both an MSSA and an MRSA strains. Interestingly, male mice challenged with an MSSA strain showed more severe and more prolonged SAGII symptomatology than animals challenged with an MRSA strain. We also showed that for male mice a high-carbohydrate diet and a high-fat diet led to asymptomatic intestinal colonization followed by delayed SAGII sign onset. In contrast, male mice fed a high-protein diet started developing mild SAGII signs early but did not develop severe SAGII until two weeks post-challenge. Furthermore, only the high-protein diet sensitized female mice to SAGII, but their symptomatology remained less severe than in male mice.

**Conclusions:** We developed a robust murine model for antibiotic-associated *S. aureus* gastrointestinal infection and colonization. This model shows both sexual dimorphism and macronutrient preference for SAGII severity. Diet manipulation can also be used to establish *S. aureus* colonization of the GI tract.

## BACKGROUND

*Staphylococcus aureus* is a gram-positive, facultative anaerobic, and opportunistic organism that causes several diseases such as skin and soft tissue infection, pneumonia, endocarditis, osteomyelitis, and bacteremia [1].

*Staphylococcus aureus* has been characterized as methicillin resistant (MRSA) or methicillin susceptible (MSSA). Although MRSA was initially described as resistant just to methicillin, these strains have acquired resistance to a broad range of antibiotics, including vancomycin, an antibiotic of last recourse against recalcitrant infections [2,3].

MRSA is often categorized into hospital-acquired (HA-MRSA) and community-acquired (CA-MRSA) infections. Similar to MRSA strains, MSSA strains can also be classified into hospital-acquired (HA-MSSA) and community-acquired (CA-MSSA). Both HA-MRSA and HA-MSSA are usually associated with surgical procedures. In contrast, CA-MRSA and CA-MSSA are mostly associated with skin and soft tissue infections [4].

*S. aureus* can deploy a veritable arsenal of distinct toxins [2]. The toxins secreted by *S. aureus* can be categorized into three groups: pore-forming toxins, exfoliative toxins (ETs), and staphylococcal enterotoxins, also known as superantigens [2]. Selective toxin production is associated with particular and unique disease presentations [5].

The mechanisms by which *S. aureus* colonizes the nares, and the skin are very well studied [6–9]. Nasal carriage is considered to be the major risk factor for surgical site infections [10]. Treatment of the nares to eliminate nasal carriage causes *S. aureus* to disappear from other body areas [11].

In contrast, the intestines have not been studied well as a potential MRSA reservoir site, even though nasal and intestinal carriage of *S. aureus* show similar levels of bacterial load [20].

Indeed, in CA-MRSA infections, *S. aureus* intestinal carriage is a prevalent risk factor [12]. These results suggest that the intestine might be a primary *S. aureus* colonization site and serve as a reservoir from which other colonization sites are replenished [13].

*Clostridioides difficile* infection (CDI) is the leading cause of antibiotic-associated diarrhea (AAD) [14,15]. Even then, CDI only accounts for approximately 25% of all events of AAD [16]. Although less prevalent than *C. difficile*, it is possible that *S. aureus* intestinal infections are underreported [17] as an etiological agent in AAD [14]. In contrast to the colonic damage caused by *C. difficile*, *S. aureus* intestinal infection seems to be established in the proximal and mid small intestines, resulting in enterocolitis and watery diarrhea [14].

Most of the studies on murine *S. aureus* are focused on skin and peritoneal infections [18,19]. A number of reports have used mice and rats as model for *S. aureus* intestinal colonization [20–24]. In contrast, *S. aureus* intestinal infection (SAGII) models have been more difficult to develop since animals develop minimal clinical signs of disease, even when *S. aureus* is inoculated at remarkably high titer [14,25,26]. To our knowledge, only one study was successful in producing severe diarrhea after direct injection of MRSA to the jejunum of immunocompromised mice [27,28]. Thus, in these models, *S. aureus* might not be truly colonizing the murine intestines but rather slowly transiting across the gut.

In this study, we developed a robust murine model for both *S. aureus* colonization and gastrointestinal infection (SAGII). We showed that antibiotic-treatment is required for the establishment of SAGII in male mice. Furthermore, we found that antibiotic-treated male mice were highly susceptible to both MSSA and MRSA strains. Interestingly, male mice challenged with an MSSA strain showed more severe SAGII symptoms and longer disease progression than animals challenged with an MRSA strain. In contrast, female mice did not develop SAGII symptoms, even when challenged with high titers of the MRSA strain. Finally, we showed that in male mice, both a high-carbohydrate diet and a high-fat diet led to asymptomatic colonization of the intestinal tract and delayed SAGII sign onset. In contrast, male mice fed a high-protein diet started developing mild SAGII signs early but eventually developed more severe disease than the other diets two weeks post-challenge. While female mice fed high-carbohydrate and high-fat diets remained resistant to SAGII infection, those fed a high-protein diet became sensitized to SAGII; however, their symptomatology remained less severe than those observed in male mice.

## MATERIALS AND METHODS

### Bacterial strains, reagents, and animal supplies

Methicillin-resistant *Staphylococcus aureus* strain FDA243 was obtained from the American Type Culture Collection (Manassas, VA). Methicillin-sensitive *S. aureus* strain RN4220 was donated by Prof. Eduardo Robleto (UNLV, Las Vegas, NV). All solid and liquid media were purchased from VWR (Radnor, PA). Antibiotics were purchased from Sigma-Aldrich (St. Louis, MO). Microbiological media and supplements were obtained from Fisher Scientific (Waltham, MA). Laboratory Rodent Standard and Experimental Diets were from LabDiet (St. Louis, MO) or Envigo (Indianapolis, IN).

### Bacterial growth conditions

The day before infection, *S. aureus* FDA243 or *S. aureus* RN4220 cells were recovered from a frozen stock and separately streaked on a Difco Tryptic Soy Agar (TSA) plate. The plate was incubated at 37 °C for approximately 14 hours to obtain individual colonies. The day of infection, a single colony of either *S. aureus* FDA243 or *S. aureus* RN4220 were individually inoculated into 15 ml of Bacto Tryptic Soy Broth (TSB) and grown at 37 °C on an orbital shaker at constant speed for approximately 6 hours to an OD of 1.5. Cells were then centrifuged for 5 minutes at 12,000 xg and resuspended into 1 ml of PBS. A 10 μl aliquot of the resulting bacterial stock was added to 90 μl of PBS and gently vortexed to mix. The bacterial suspension was then serially diluted from 10 ¹ to 10 of the original stock.

Each dilution was spot-plated on a TSA plate and incubated at 37 °C overnight to determine bacterial titer. We consistently obtained a final concentration of approximately 10^10^ CFUs per ml stock solution. The stock solution was diluted as needed to obtain the appropriate inocula for animal experiments.

### Animals

Animal protocols were performed in accordance with the Guide for Care and Use of Laboratory Animals outlined by the National Institutes of Health. Protocols were reviewed and approved by the Institutional Animal Care and Use Committee (IACUC) at the University of Nevada, Las Vegas (Permit Number: R0914-297). Weaned male and female C57BL/6J mice were obtained from Jackson Labs, Jax west (Bar Harbor, ME). Mice were housed in groups of five mice per cage at the University of Nevada, Las Vegas animal care facility. Upon arrival at the facility, mice were quarantined and allowed to acclimate for at least one week prior to experimentation. All post-challenge manipulations were performed within a biosafety level 2 laminar flow hood.

### Developing the murine model of antibiotic-associated *S. aureus* intestinal infection (SAGII)

The murine SAGII model developed in this study was adapted from the CDI mouse model [15]. To test for the effect of antibiotic regime on SAGII, groups of male mice were dosed for 0 (n=5), 7 (n=5), or 14 (n=10) consecutive days of an *ad libitum* aqueous solution of an antibiotics cocktail consisting of vancomycin (0.045 mg/ml), metronidazole (0.215 mg/ml), gentamicin (0.035 mg/ml), colistin (850 U/ml), and kanamycin (0.4 mg/ml). Mice were then switched to autoclaved deionized (DI) water for the rest of the study. Two days prior to infection (day-2), mice were administered an intraperitoneal (IP) injection of 10 mg/kg clindamycin. On the day of infection (day 0), animals were challenged with 10^9^ CFUs of *S. aureus* strain FDA243 by oral gavage.

To test for the effect of *S. aureus* titer on SAGII, groups of male mice were treated as above, but were challenged with either 10^7^ (n=5), 10^4^ (n=10), or 10^2^ (n=10) CFUs of *S. aureus* strain FDA243.

To test for sexual dimorphism in SAGII, groups of female mice were treated as above and challenged with either 10^9^ (n=10), 10^7^ (n=10), 10^4^ (n=5), or 10^2^ (n=5) CFUs of *S. aureus* strain FDA243.

To test for the effect of diet on SAGII, groups of male (n=5 or n=10 per diet) and female (n=5 per diet) mice were fed either a high-protein diet (59%P, 20.7%C, 20.2%F), a high-carbohydrate diet (5%P, 75.7%C, 19.3%F), or a high-fat diet (5.1%P, 22.1%C, 72.8%F) for 10 days prior to antibiotic treatment. While still on their corresponding diets, mice were treated with antibiotics, as above. On day 0, mice were challenged with 10^2^ CFUs of *S. aureus* strain FDA243.

To test for *S. aureus* strain effects on SAGII, male mice were treated as above and challenged with either 10^2^ CFUs of methicillin-resistant *S. aureus* strain FDA243 (n=10) or methicillin-sensitive *S. aureus* strain RN4220 (n=5).

In all experiments, mice were observed for signs of infection daily and disease severity was scored according to a sign rubric adapted from the murine CDI model [29]. According to the rubric, pink anogenital area, mild wet tail, and weight loss of 8-15% were each given an individual score of 1. Red anogenital area, lethargy/distress, increased diarrhea/soiled bedding, hunched posture, and weight loss greater than 15% were each given an individual score of 2.

After all scores were summed, animals scoring less than 3 were considered non-diseased and were indistinguishable from non-infected controls. Animals scoring 3–4 were considered to have mild SAGII. Animals scoring 5–6 were considered to have moderate SAGII. Animals scoring greater than 6 were considered to have severe SAGII and were immediately culled.

### Statistical analysis

Murine SAGII sign severity was analyzed via box-and-whisker plots with a minimum of five independent values (n ≥ 5). A single factor ANOVA or one-tailed *t*-test was performed using the JASP program to assess differences between groups at every time point. ANOVA results with *p*-values <0.05 were analyzed *post hoc* using the Scheffé test for comparison between all groups.

## RESULTS

### Effect of antibiotics regime on gastrointestinal S. aureus infection

To establish a murine model of *S. aureus* antibiotic-associated gastrointestinal infection (SAGII), we treated male mice with antibiotics for 0, 7, or 14 days prior to challenge with 10^9^ CFUs of MRSA strain FDA243. Antibiotic-treated animals started to develop gastrointestinal symptoms (soft stool, mild wet tail, pink anogenital area, and weight loss) by day 1 post-challenge. For both antibiotic-treated groups, sign severity continued to increase (severe diarrhea, severe wet tail, red anogenital area, weight loss, lethargy, and death) and reached a maximum by days 2-3 post-challenge. SAGII signs started to subside by day 4 post-challenge and most surviving animals became asymptomatic by days 6 (**Figure 1**).

**Figure 1.**
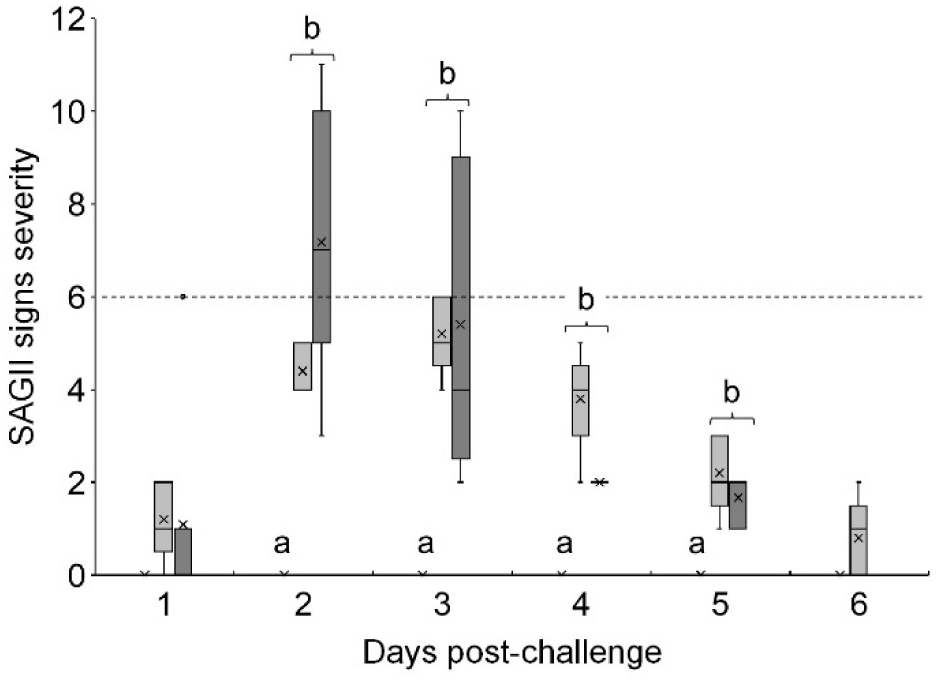
The effect of antibiotic duration on *S. aureus* infection: Box-and Whisker-plots of SAGII signs severity in male mice treated with different antibiotic regimes and challenged with *S. aureus* strain FDA243. Mice were treated with either water (white bars), antibiotic cocktail for 7 days (light gray bars), or antibiotic cocktail for 14 days (dark gray bars) before challenge with 10^9^ CFUs of MRSA strain FDA243. SAGII signs severity were determined based on the CDI scoring rubric reported previously [29]. Animals with a score of <3 were indistinguishable from non-infected controls and were considered non diseased. Animals with a score of 3 to 4 were considered to have mild SAGII. Animals with a score of 5 to 6 were considered to have moderate SAGII. Animals with a score of >6 (dashed line) were considered to have severe SAGII and were immediately culled. Horizontal bold lines represent median values, asterisks represent mean values, bars represent interquartile ranges, whiskers represent maximum and minimum values, and dots represent inner point values. Single-factor ANOVA was performed at every time point to assess differences among the sign severity means of the untreated, 7-day antibiotic treated, and 14-day antibiotic treated groups. Columns that are labeled with different letters are statistically different (*P* > 0.05) by one-way ANOVA followed by *post hoc* Scheffé test for comparison between all groups.

Animals on the 14-day antibiotic regimen showed more individual sign variability compared with animals on the 7-day regimen. Some animals on the 14-day antibiotic regimen became deathly ill from SAGII, while others showed less symptoms than animals on the 7-day regime. Because of these variabilities, there was no statistical difference in SAGII sign severity between the two antibiotic treatment groups. Non-antibiotic treated animals never developed any signs of infection.

### Effect of bacterial load on gastrointestinal S. aureus infection

To test the effect of bacterial load on infection severity, we treated male mice with a 7-day antibiotic course prior to challenge with either 10^2^, 10^4^, 10^7^, or 10^9^ CFUs of MRSA strain FDA243. SAGII animals challenged with lower bacterial loads still developed signs comparable with animals challenged with 10^9^ CFUs of MRSA strain FDA243 (**Figure 2A**), even though the course of the infection was delayed by approximately 24 hours compared to animals challenged with the highest bacterial load.

**Figure 2.**
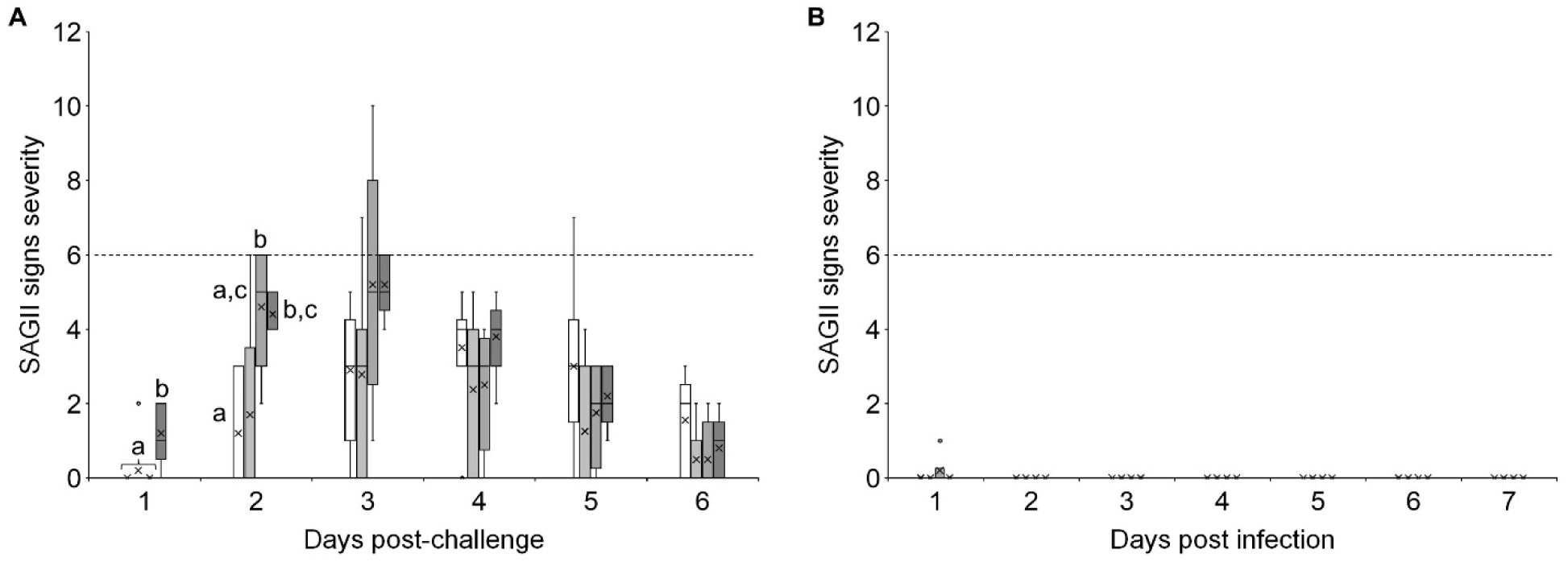
Determination of lowest dosage to induce clinical symptoms: Box-and-Whisker plot of SAGII signs severity in (A) male mice and (B) female mice challenged with different titers of *S. aureus* strain FDA243. Four groups of male and four groups of female mice were treated with antibiotics for 7 days prior to infection. Mice were then infected with 10^2^ CFUs (white bars), 10^4^ CFUs (light grey bars), 10^7^ CFUs (medium grey bars), or 10^9^ CFUs (dark grey bars). SAGII signs severity and statistical analysis were determined as above. Columns that are labeled with different letters are statistically different (*P* > 0.05) by one-way ANOVA followed by *post hoc* Scheffé test for comparison between all groups.

### Effect of sex on gastrointestinal S. aureus infection

To test the effect of sex on SAGII, we treated female mice with a 7-day antibiotic course prior to challenge with either 10^2^, 10^4^, 10^7^, or 10^9^ CFUs of MRSA strain FDA243. In contrast to male mice, none of the female mice developed infection signs, even at the highest bacterial load tested (**Figure 2B**).

### The effect of diet on gastrointestinal S. aureus infection

To test the effect of diets on infection severity, individual groups of male and female mice were fed either a high-carbohydrate, high-fat, or high-protein diet for 14 days prior to challenge with 10^2^ CFUs of MRSA strain FDA243. In contrast to the rapid infection onset observed for male mice fed a standard diet (**Figure 2A**, white bars), animals fed a high-fat and high-carbohydrate diet did not develop symptoms for at least 10 days post-challenge (**Figure 3A**). Even though SAGII onset was delayed, sign severities for animals that were fed a high-carbohydrate or high-fat diet was similar to the animals that were fed the standard diet.

**Figure 3.**
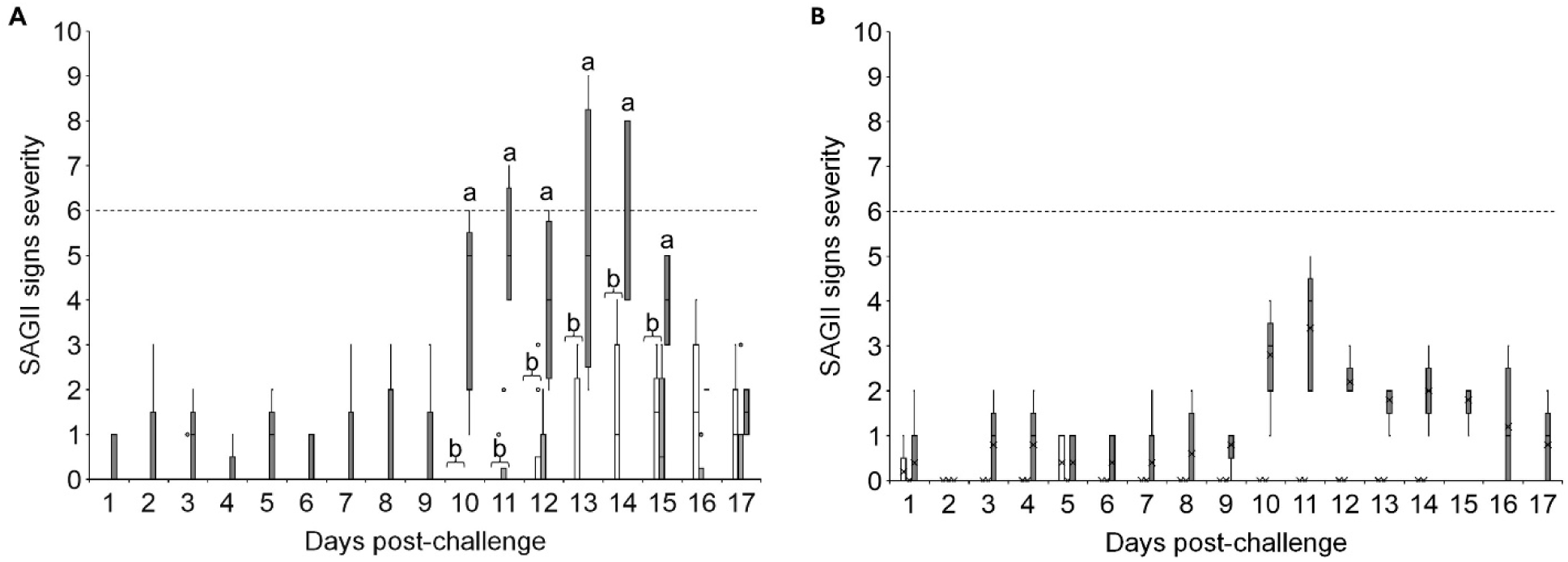
The effect of diet on *S. aureus* infection in male mice: Box-and-Whisker plot of SAGII signs severity in (A) male mice and (B) female mice fed with different diets and challenged with *S. aureus* strain FDA243. Three groups of male and three groups of female mice were fed a high-carbohydrate diet (white bars), a high-fat diet (light gray bars), or a high-protein diet (dark gray bars). Animals were then treated with antibiotics for 7 days prior to challenge with 10^2^ CFUs of *S. aureus* strain FDA243. SAGII signs severity and statistical analysis were determined as above. Columns that are labeled with different letters are statistically different (*P* > 0.05) by one-way ANOVA followed by *post hoc* Scheffé test for comparison between all groups.

In contrast, animals that were fed a high-protein diet started developing signs on day 1 post-challenge and continued showing mild SAGII signs for 9-days post-challenge. By day 10, however, animals in the high-protein diet group started to develop moderate to severe SAGII signs that were statistically higher than for animals in any other diet. Severe SAGII signs persisted for at least five more days, with most animals becoming deathly ill.

Female mice fed the high-carbohydrate, or the high-fat diet were as refractive to MRSA challenge as animals in the standard diet and barely developed any signs of infection (**Figure 3B**). In contrast, female mice fed a high-protein diet did develop SAGII signs with similar infection onset delay as their male counterpart, albeit the infection was less severe than in male mice fed the same diet.

### The effect of S. aureus strains on gastrointestinal infection

To test the effect of different *S. aureus* strains on infection severity, we treated male mice with a 7-day antibiotic course prior to challenge with 10^2^ CFUs of either MRSA strain FDA243 or MSSA strain RN4220. Both strains caused similar infection progression with animals starting to show SAGII signs by day 2 post-challenge and infection severity peaking at around day 3 (**Figure 4**). However, by day 3 post-challenge, MSSA-challenged animals showed statistically more severe disease than MRSA-challenged animals. Furthermore, while animals infected with the MRSA strain completely recovered by day 7 post-challenge, animals infected with the MSSA strain remained symptomatic for at least three more days.

**Figure 4.**
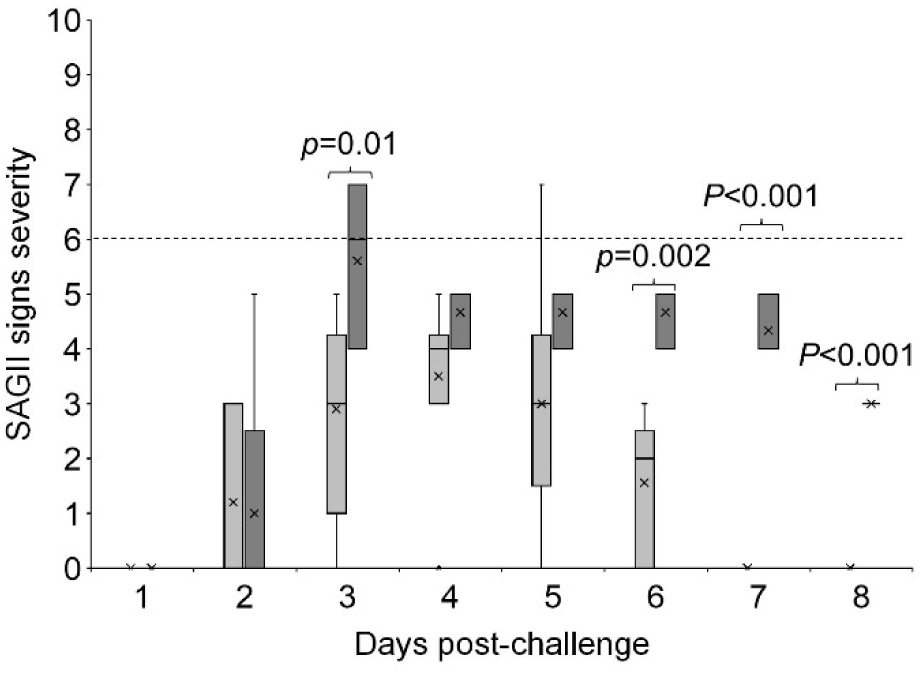
The effect of *S. aureus* MSSA strain in gastrointestinal infection. Box-and Whisker-plot of SAGII signs severity in male mice challenged with MRSA strain FDA243 (light gray bars) or MSSA strain RN4220 (dark gray bars). Two groups of male mice were treated with antibiotics for 7 days prior to challenge with 10^2^ CFUs of *S. aureus* strain FDA243 or *S. aureus* strain RN4220. SAGII signs severity and statistical analysis were determined as above. Daily statistical differences between groups were determined by Student *t*-test. P-values are shown for statistical differences (*P* < 0.05) between the two conditions.

## DISCUSSION

Even though *C. difficile* is the main identifiable agent of antibiotic-associated diarrhea (AAD), it is still only responsible for 15-25% of cases [16]. Although less studied, *S. aureus* can also cause AAD [14,30]. Previous efforts to create a mouse model for *S. aureus* intestinal infections showed that animals could be colonized by *S. aureus* for weeks post-challenge [14]. However, *S. aureus* infection in these models did not appear to induce clinical signs of disease.

To develop a *S. aureus* infection model, we followed our successful antibiotic treatment regimen developed for the murine model of CDI [15]. Male mice seemed to be very susceptible to *S. aureus* infection. Indeed, animals developed moderate to severe signs, regardless of the bacterial load used for challenge. Remarkably, even 100 bacterial cells were sufficient to cause symptoms of intestinal infection similar to animals challenged with a 10-million times higher infective dose.

The proportion of macronutrients in a diet has been shown to affect CDI severity and outcome in both mice and hamsters [31,32]. We saw a complex diet effect for SAGII onset. When male mice were put on high-carbohydrate or high-fat diets, infection onset was delayed for over a week compared to the standard diet. However, the severity of symptoms for delayed-onset SAGII (high-carbohydrate and high-fat diets) was not different than symptoms in early-onset SAGII (standard diet). In contrast, animals in a high-protein diet started to show SAGII signs early and eventually developed more severe signs than animals fed any other diet.

In contrast to their male counterparts, female mice were very resistant to *S. aureus* intestinal infections. Furthermore, diet composition had a minor effect on the susceptibility of female mice to SAGII. Neither the standard diet, high-carbohydrate diet, nor the high-fat diet were able to induce infection signs in female mice. Only animals in high-protein diet showed delay-onset of SAGII, but even then, female mice developed milder symptoms than males. The severity of infection varied among the two *S. aureus* strains tested. Indeed, the MSSA strain showed to produce more severe and deadly disease than the MRSA strain.

Furthermore, while surviving MRSA-infected animals recovered by day 6 post-challenge, surviving MSSA-infected animals still showed statistically more severe symptoms even 8 days post-challenge.

## CONCLUSIONS

By adapting our murine model of CDI, we were able to build a robust murine model for *S. aureus* gastrointestinal infection (SAGII) and colonization. We showed that dysbiosis of the gut microbiome following antibiotics usage needs to occur for *S. aureus* to be able to colonize in the gut and establish infection. To our knowledge, this is the first report of a murine model of *S. aureus* intestinal infection where animals develop significant clinical signs.

The fact that MSSA strain RN4220 caused more severe symptoms than MRSA strain FDA243 is intriguing. These differences could be attributed in part to differential toxin production between the two strains. Indeed, it has been reported that MRSA strains evolved from MSSA strains when they acquired the Staphylococcal Cassette Chromosome mec (SCCmec) elements that play a major role in antibiotics resistance [33,34]. Interestingly, some HA-MRSA strains containing the SCCmec type II element were less virulent than MSSA strains [35]. Furthermore, the *mecA* gene which is responsible for β-lactam antibiotic resistance, is associated with reduced *S. aureus*-mediated toxicity [36].

The virulence factors of laboratory MSSA strain RN4220 used in this study has been well characterized. MSSA strain RN4220 does not secrete superantigens nor large amounts of cytotoxins [37]. Instead, MSSA strain RN4220 secretes β-toxin in large quantities targeting sphingomyelin in host cell membranes, leading to lysis [38].

MRSA strain FDA243 was isolated from the fecal samples of a sick child with *S. aureus* gastrointestinal infection and is less well characterized. It is, however, reported that MRSA strain FDA243 produces Staphylococcal enterotoxin B (SEB), which is believed to be the primary toxin responsible for inflammation of the small intestine during *S. aureus* gastrointestinal infections [14]. Interestingly, SEB did not seem to be required for tissue damage in mouse models, suggesting that other virulence factors could be associated with damage in the small intestines following an infection [14].

We found a clear sexual dimorphism for SAGII, with male mice being very susceptible to infection and females were quite resistant. This is not unexpected since bacterial infections have known sexual dimorphism [26] with females tending to be more susceptible to genitourinary tract infections [27, 28], while males are more prone to gastrointestinal [29] and respiratory infections [30]. Indeed, human males have an increased risk of developing bacteremia or bloodstream infection with both MSSA and MRSA compared to human females [39].

A recent study found similar sexual dimorphism for murine intestinal colonization by *S. aureus*. This study showed that differences in microbiota and immune response of female mice compared to male mice were key players in their protection against MRSA gastrointestinal colonization [40]. Whether the same mechanism also protect female mice from *S. aureus* gastrointestinal infection needs to be established. Similarly, it would be interesting to determine if female SAGII protection is correlated with estrous cycling and concomitant sex hormone changes.

Intriguingly, we have observed the opposite type of sexual dimorphism in murine CDI. Indeed, we recently showed that female mice developed more severe CDI than their male counterparts [41]. Furthermore, we found that CDI sign severity in female mice correlated with the estrous stage and negatively correlated with the serum levels of sexual hormones necessary for estrous cycling [41]. It will be interesting to determine the effect of hormone levels on the protection of female mice against SAGII and contrast them with the sexual dimorphism effect seem in CDI [38].

We also found macronutrient effects on SAGII virulence. High-carbohydrate and high-fat diets both delay SAGII onset in male mice but do not affect the severity of the infection. We attribute this delay to the ability of *S. aureus* to colonize the murine intestine. However, it is not clear what triggers the established *S. aureus* cells to initiate the delayed infection.

Prior work on the effect of diets on *S. aureus*-induced murine sepsis showed that polyunsaturated high-fat or low-fat diets resulted in higher survival rate and decreased renal bacterial loads compared to a saturated high-fat diet [42]. These contrasting effects of dietary fat on *S. aureus* infection outcomes could be due to multiple factors, including different infected tissues, bacterial strains, antibiotic interventions, and/or challenge administration.

In contrast to the delayed effect of the high-carbohydrate and high-fat diets on SAGII, neither diet affects murine CDI onset [43]. Furthermore, high-carbohydrate diets tend to be protective against CDI, while high-fat diets can increase CDI severity [43]. The differential effects of these diets on SAGII and CDI is intriguing. These divergences might be due to diet-induced changes of the murine microbiome [43] and the unique way that they affect each infective agent.

While a high-protein diet does not delay the initial infection sign development, it exacerbates SAGII in male mice and, intriguingly, also in female mice. We have previously seen a similar trend in the murine CDI model. Indeed, an Atkins-type diet (high-fat/high-protein) did not cause delay in infection onset, but greatly increased CDI severity [43].

In conclusion, we have developed a mouse model for both *S. aureus* intestinal infection and colonization. By varying mouse sex and dietary macronutrients, this new murine SAGII model can be controlled in terms of both timing of disease onset and sign severity.

## Acknowledgements

This work was supported by the National Institute of Allergy and Infectious Diseases at the National Institute of Health [grants numbers R15AI164346, R16AI175022] to EAS, and by the National Institute of General Medicinal Sciences at the National Institute of Health [grant number P20GM12132] to NIPM. We would also like to extend our gratitude to Dr. Chandler Hassan, Rose Jiang, Naomi Okada, and Erica Bacab for their assistance and support throughout this study. Lastly, we would like to thank the mice for their contribution to science.

## Author Contributions

EAS conceived the development of the murine *S. aureus* gastrointestinal infection model and supervised the project. MN established the mouse SAGII model and directed all the experiments described. SY, LB, LW, and EH performed the experiments described. EAS and MN designed experiments, analyzed data, and drafted the manuscript.

